# The dynamic linkage between intact provirus integration sites and the host functional genome property alongside HIV-1 infections associated with antiretroviral therapy

**DOI:** 10.1101/2022.12.02.518849

**Authors:** Heng-Chang Chen

## Abstract

The HIV-1 latent reservoir harboring replication-competent proviruses, is the major barrier in the quest for a HIV-1 infection cure. HIV-1 infection at all stages of disease progression is associated with immune activation and dysfunctional production of proinflammatory soluble factors (cytokines and chemokines) and it is expected that during HIV-1 infection different immune components and immune cells, in turn, participate in immune responses, subsequently activating downstream biological pathways. However, whether HIV-1 infections activate only specific pathways or result in the global activation of functional pathways is presently not fully understood. Therefore, in this work, I used genes targeted by intact proviruses from published datasets to seek enriched immunologic signatures and host biological pathways alongside HIV-1 infections. I observed that different compositions of immune cell types and proinflammatory soluble factors appeared alongside HIV-1 infections associated with antiretroviral therapy based on the over-representation analysis. Moreover, KEGG pathways relevant to “cancer specific type”, “immune system”, “infectious disease viral” and “signal transduction” were frequently enriched in HIV-1-infected individuals subjected to antiretroviral therapy.

## Introduction

HIV-1 infection is a dynamic disease progression associated with a combination of immune cells, immune components, and various cellular restriction factors aimed at battling infections. It is known that HIV-1 infection induces a plethora of proinflammatory soluble factors (cytokines and chemokines), that recruit and activate innate immune-related cells to the site of HIV infection to restrict the replication and spread of viruses. Even though this defense mechanism initiates at the early stage of HIV-1 infections, HIV still manages to complete its life cycle and establish latent reservoirs.

Several host genes, so-called recurrent integration genes (Ikeda et al., 2007; Maldarelli et al., 2014; Wagner et al., 2014) were demonstrated to be frequently targeted by HIV-1 in HIV-1-infected individuals. However, the mechanism that leads HIV-1 integration sites to be present in these recurrent integration genes and the biological meaning of this phenomenon remains unclear. Nevertheless, a remarkable study attempted to answer this question by showing the interaction between the selection of HIV integration sites and functional pathways assigned by different biological processes in the host cells (Zhyvoloup et al., 2017). Their findings implied that the possible mechanism leading to HIV-1 integration in hot spot genes can happen at the level of host functional property formed by a group of gene sets rather than a gene per se.

In the continuation of this concept, in this work, I sought any dynamic linkage between the configuration of the HIV-1 reservoir and host immunity alongside HIV-1 infections. I used the host genes targeted by intact proviruses (Einkauf et al., 2022; Jiang et al., 2020) as an input gene list because only intact proviruses are responsible for viral rebounds. This list of genes included integration sites retrieved from elite controllers (Jiang et al., 2020) and HIV-1-infected individuals subjected to antiretroviral therapy (ART) (Einkauf et al., 2022). Of note, integration sites published by Einkauf et al. (2022) (Einkauf et al., 2022) were longitudinally collected from five HIV-1-infected individuals, thereby allowing us to follow how the HIV reservoir evolves during the period of ART. Based on the over-representation analysis, I observed that different immunologic signatures were enriched alongside HIV-1 infections associated with ART. In contrast, few immunologic signatures were enriched in elite controllers. In addition, enriched gene sets were mainly involved in biological pathways related to “cancer specific type”, “immune system”, “infectious disease viral” and “signal transduction”. Finally, I found that enriched immunological signatures were contributed by intact proviruses-targeted genes that are associated with gene products of HIV-1 or affect HIV-1 replication and infectivity. Based on these findings, I proposed a hypothesis that there could be a dynamic interplay between HIV-1 integration sites and the host functional genome property: HIV-1 integration frequency might be used as a surrogate for gene sets, which may define specific immune cell types and proinflammatory soluble factors during HIV-1 infection.

## Results

### Different immunologic signatures were enriched alongside HIV-1 infections associated with ART

121, 149, and 176 genes from pretreatment HIV-1-infected individuals (ul), HIV-1-infected individuals subjected to a short-(st) and a long period of ART (lt) (Einkauf et al., 2022) (Table 1), and 104 genes from elite controllers (Jiang et al., 2020) (Table 1) were employed to perform over-representation analysis, respectively.

**Table 1.**
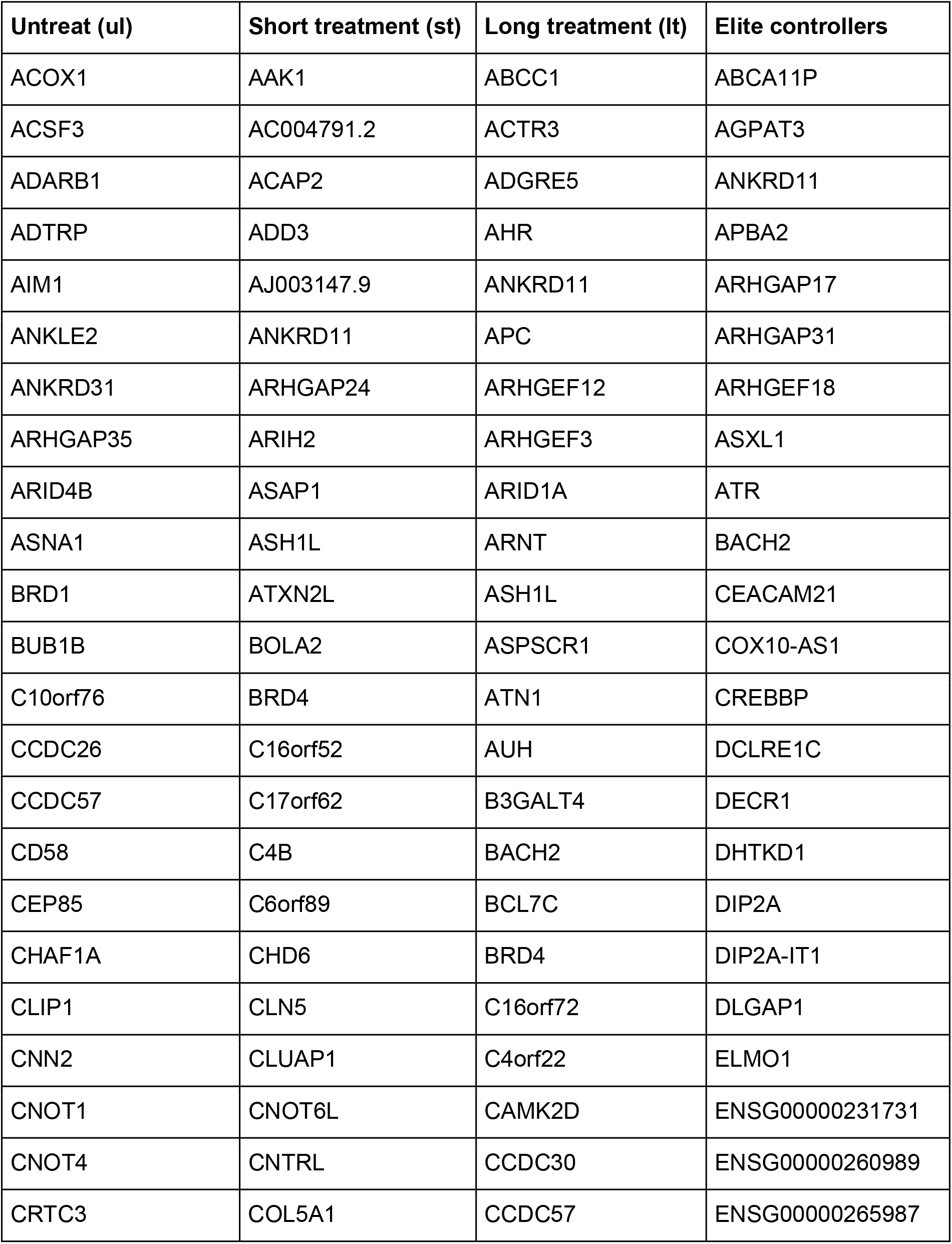
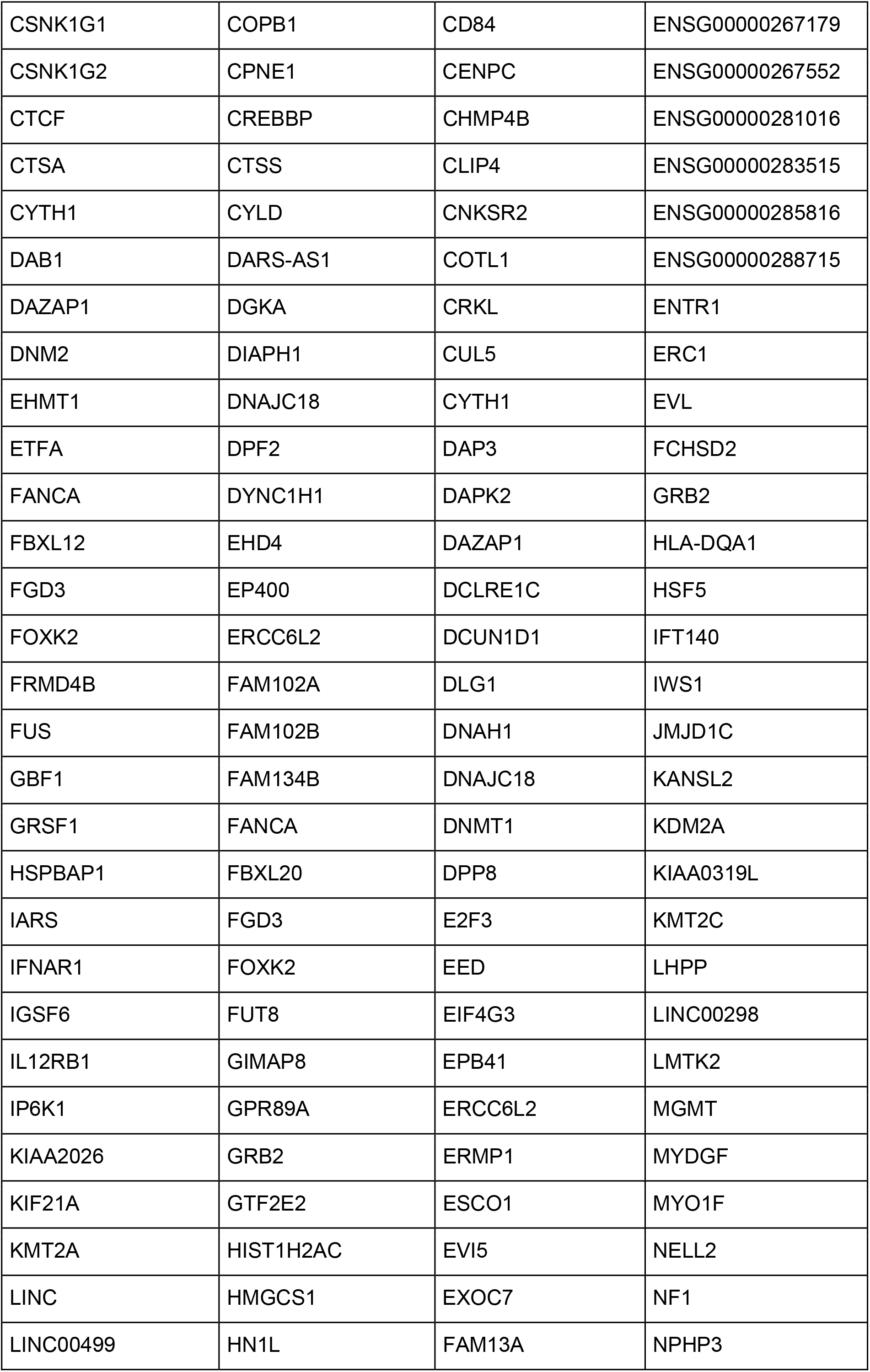
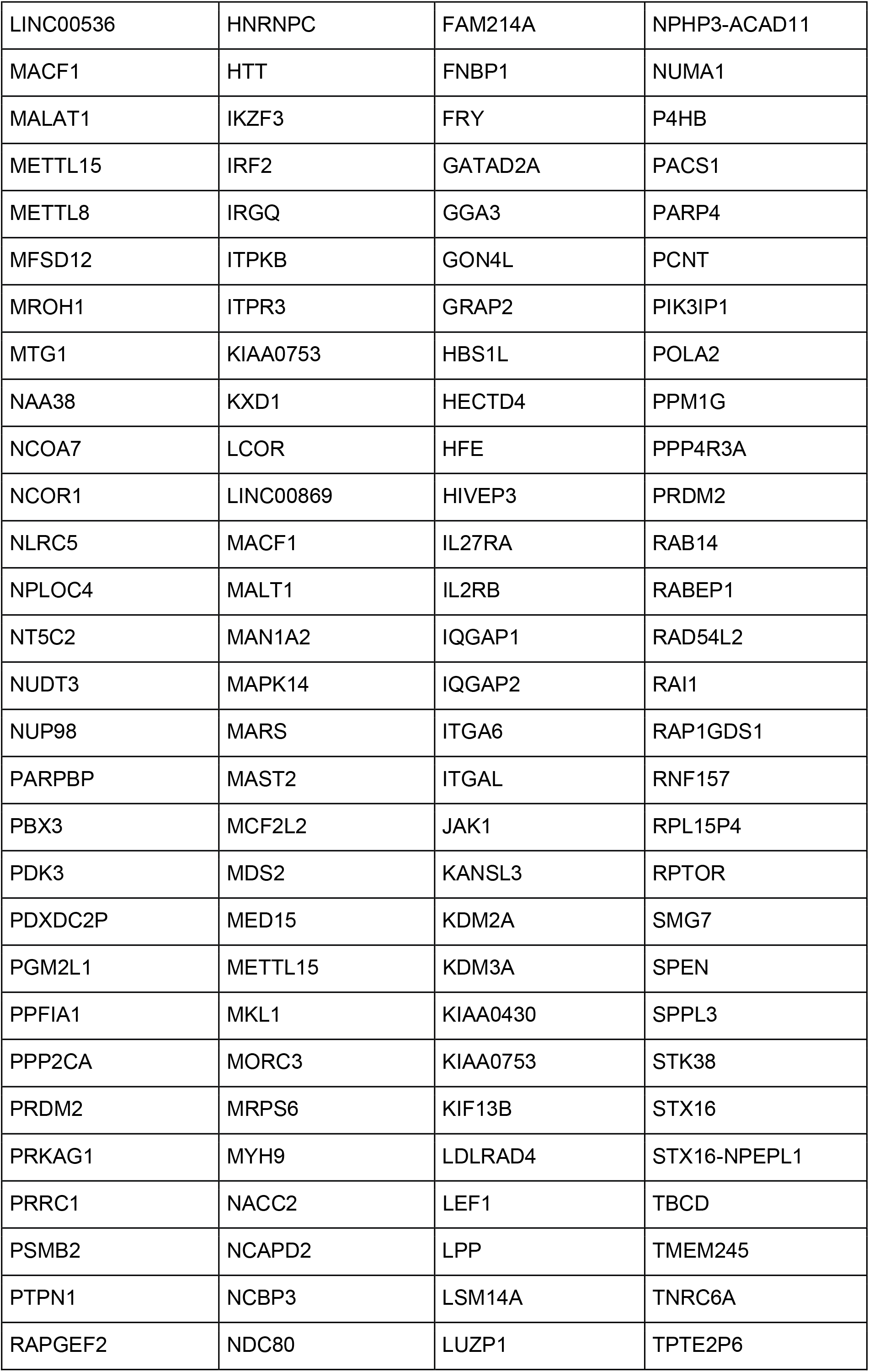
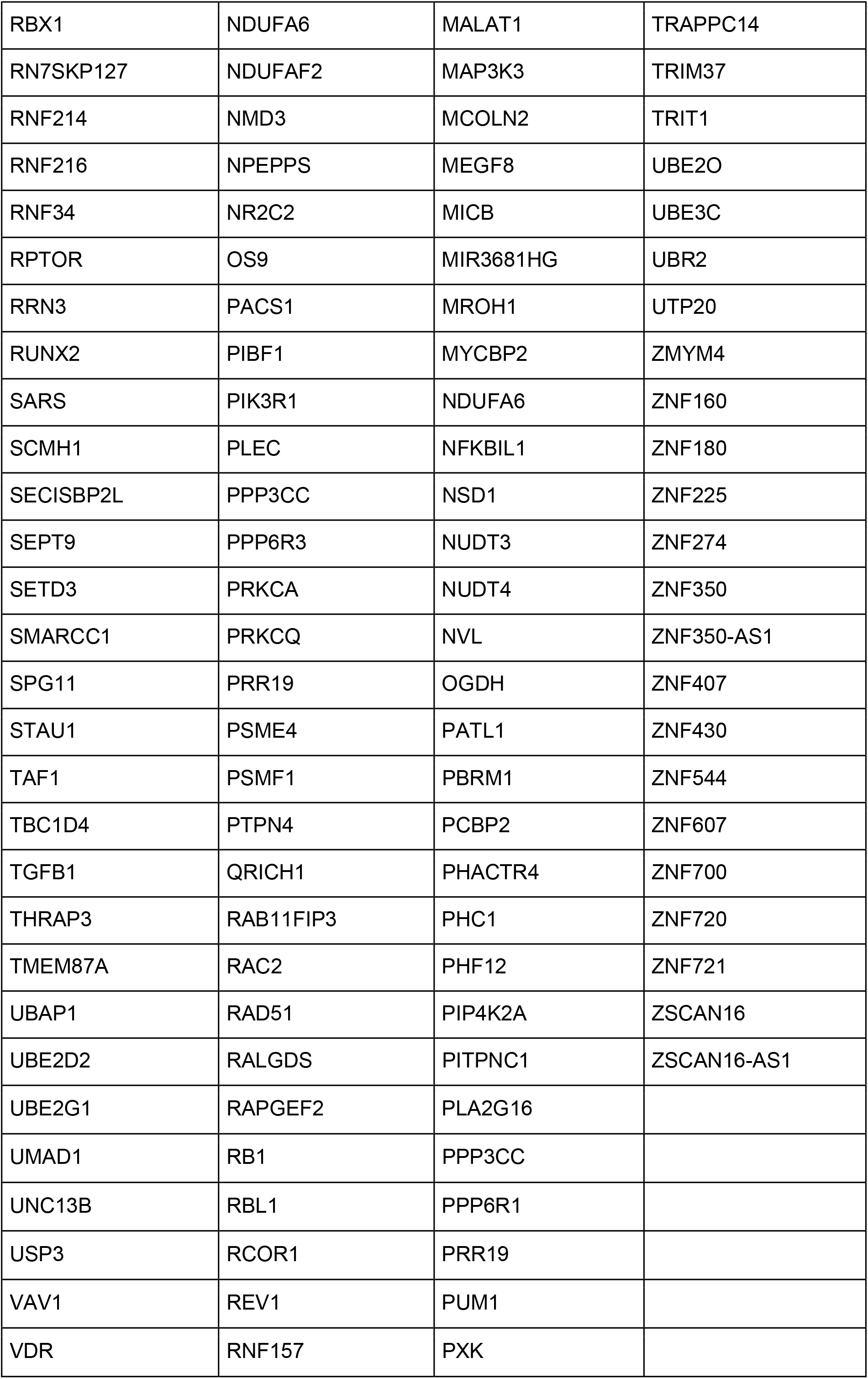
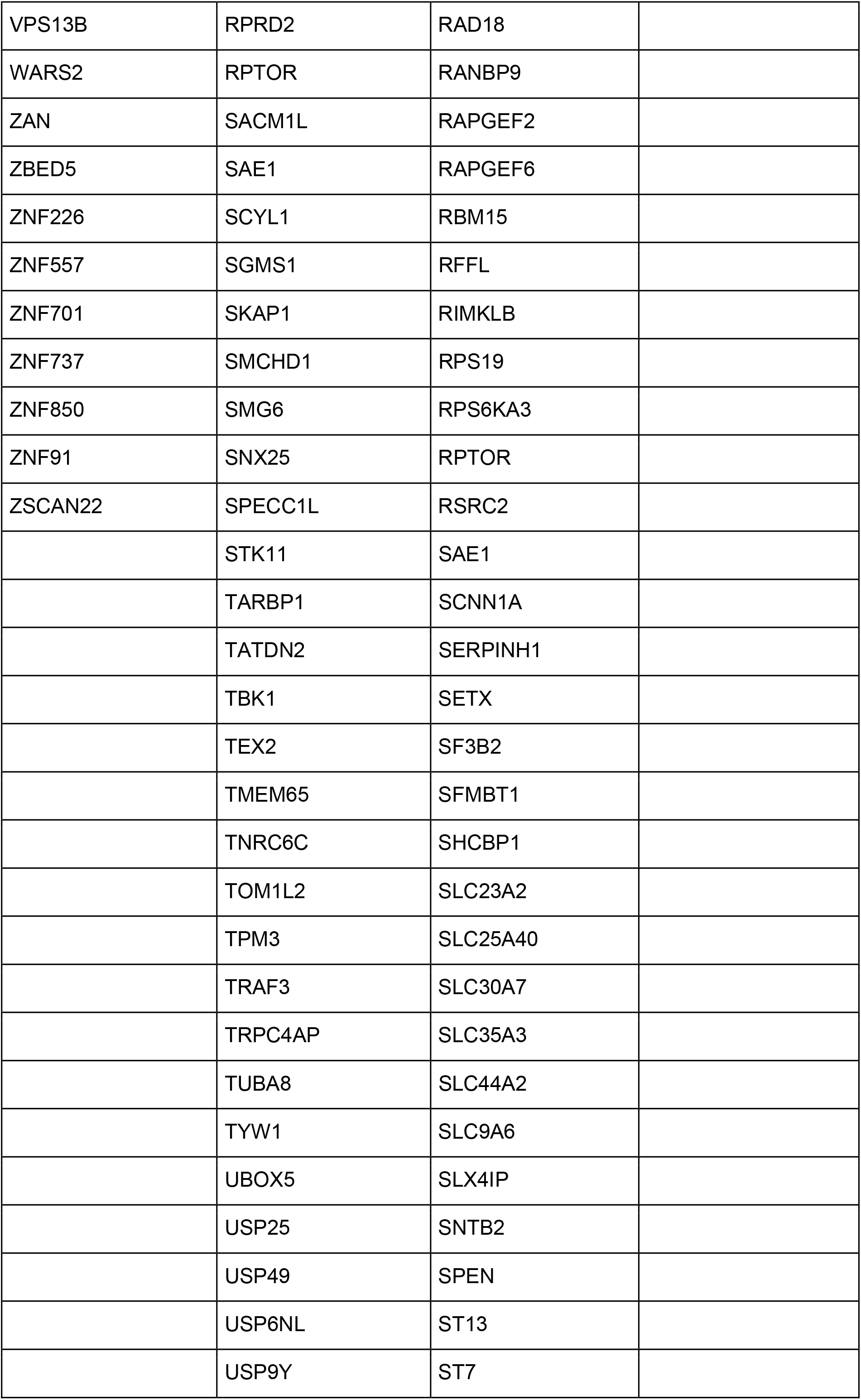
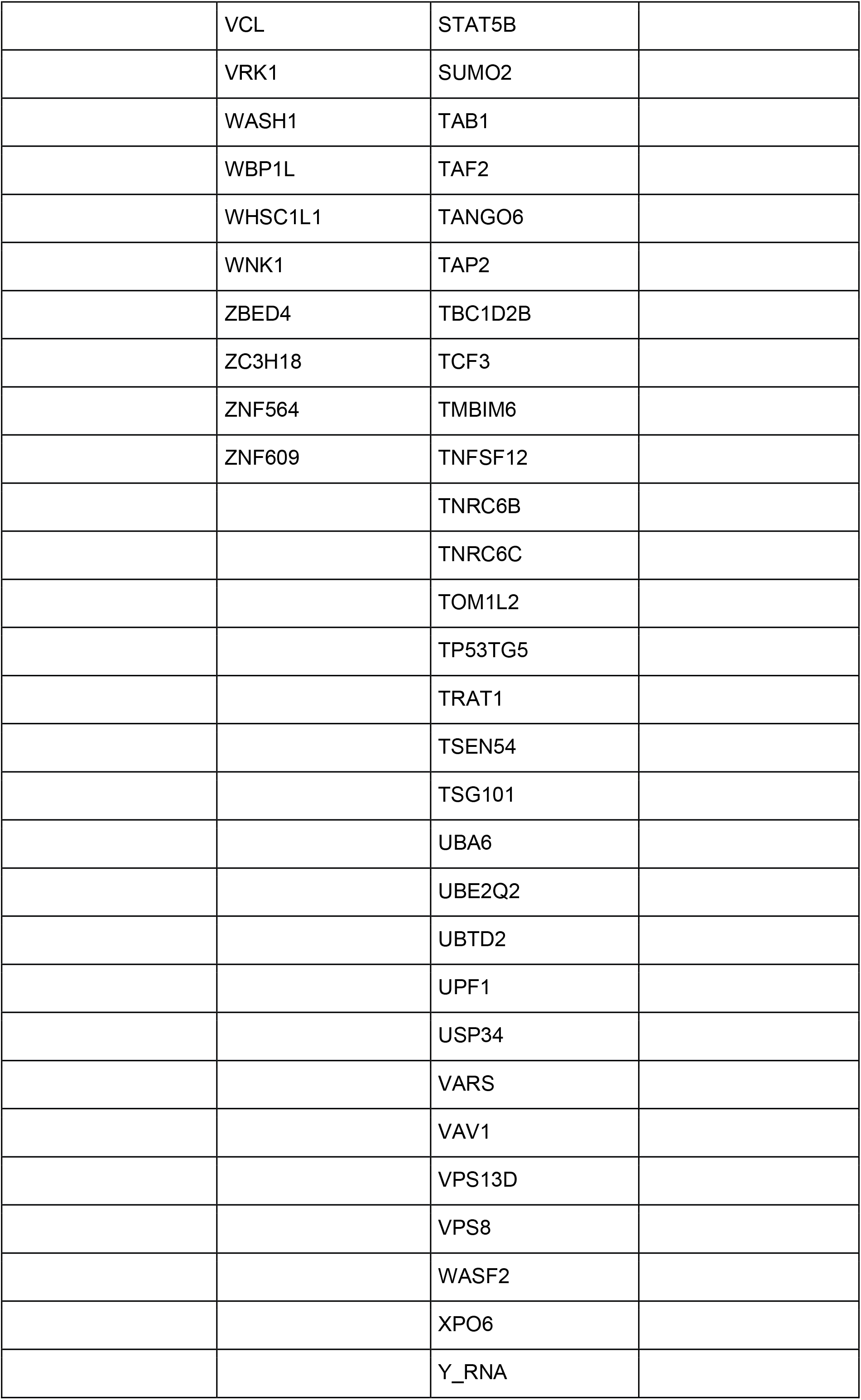
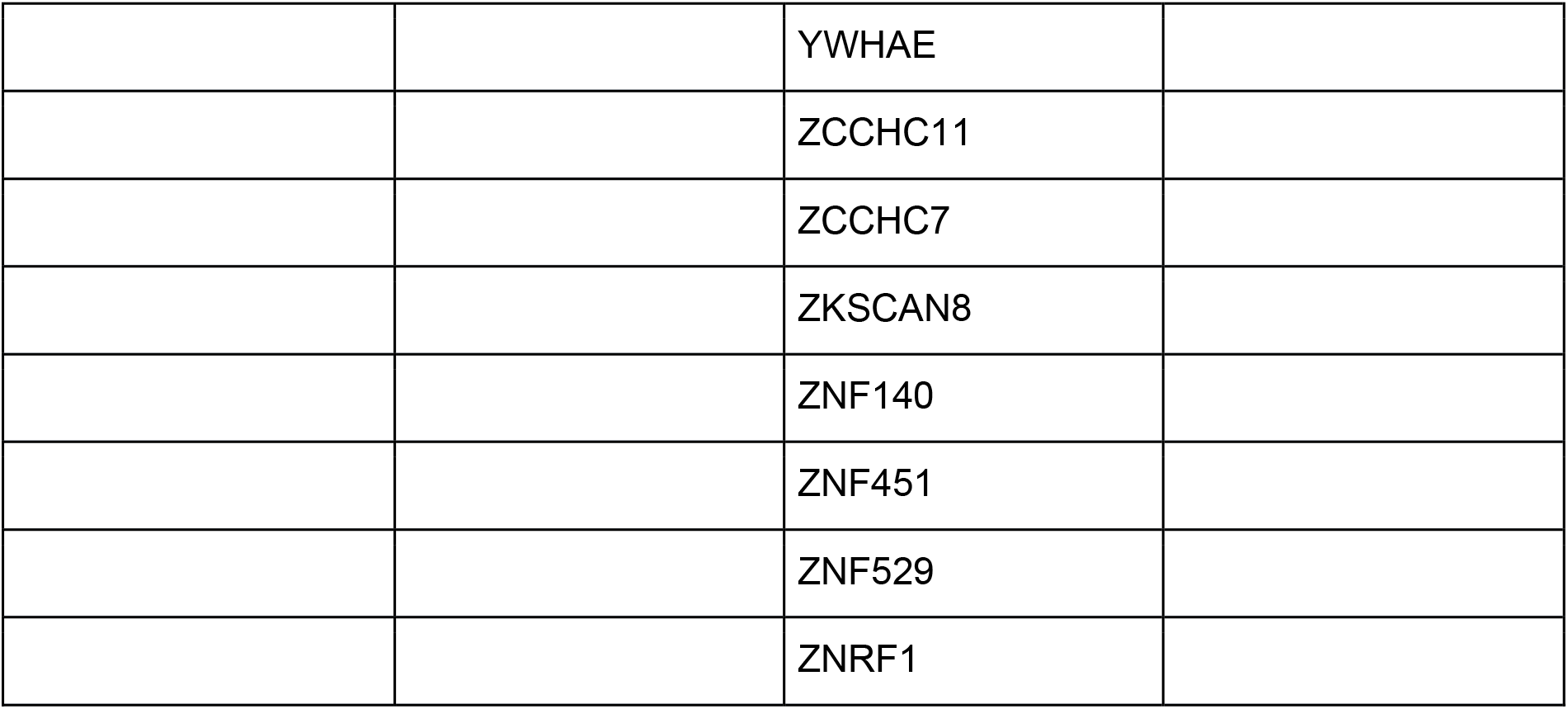
Input gene list

A list of the intact-provirus-targeted genes used for over-representation analysis in this work. Intact provirus integration sites were retrieved from Jiang et al. (2020) (Jiang et al., 2020) for elite controllers and Einkauf et al. (2022) (Einkauf et al., 2022) for HIV-1-infected individuals.

8 (S1 Table), 65 (S2 Table), 152 (S3 Table), and 22 (S4 Table) immunologic signatures were enriched in pretreatment HIV-1-infected individuals and HIV-1-infected individuals subjected to a short- and a long period of ART and in elite controllers, respectively (Fig 1A). It is essential to stress that due to a lack of expression data corresponding to integration sites used in this work, I, therefore, calculated rich factors [21] (see Materials and Methods) rather than net enrichment scores, to represent the enrichment intensity. I used randomly selected 150, 250, 350, 450, 650, and 1000 genes (data not shown) and the whole genome to perform the over-representation analysis as controls to validate the significance of rich factors obtained in HIV-1-infected individuals and elite controllers (Fig 1B). None of the immunologic signatures was enriched using 150, 250, 350, 450, 650, and 1000 genes (data not shown), and one immunologic signature was enriched using randomly selected 550 genes (rich factor: 3.681). Although 4872 immunological signatures were enriched using the whole genome (rich factor: median, 1.162; mean 1.154), rich factors measured in HIV-1-infected individuals (both pretreatment and receiving ART) and in elite controllers showed significance compared to controls (Fig 1B), indicating that immunological signature gene sets detected in this study were significantly enriched.

**Fig 1.**
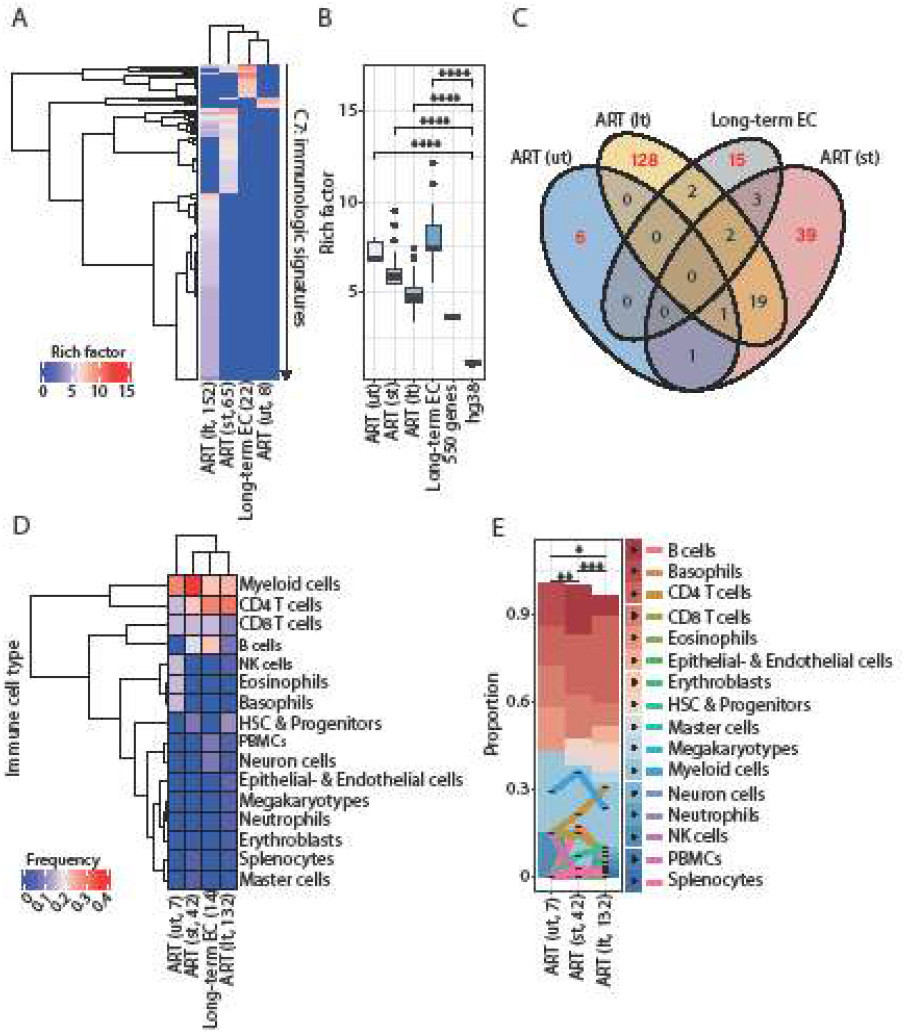
Distinct immunologic signatures were enriched in HIV-1-infected individuals and elite controllers. (A) Cluster heatmap representing immunologic signatures enriched in HIV-1-infected individuals and elite controllers. Parenthesis placed after the column name indicates the status of the antiretroviral therapy (“lt” is referred to as HIV-1-infected individuals with long-term ART; “st” is referred to as HIV-1-infected individuals with short ART; “ut” is referred to as pretreatment HIV-1-infected individuals) and the quantity of enriched immunologic signatures. The color scale represents the magnitude of the enrichment represented by rich factors. (B) Box plots representing the magnitude of enrichment of immunological signatures. Commands executed to calculate rich factors were described in Materials and Methods. Statistic significance was calculated by the Wilcoxon test by R with default options. (C) Venn diagram representing the overlap of enriched immunologic signatures in HIV-1-infected individuals and elite controllers. Red numbers indicate the quantity of unique immunologic signatures enriched in pretreatment HIV-1-infected individuals, HIV-1-infected individuals subjected to ART and elite controllers. (D) Cluster heatmap represents the frequency of immune cell types enriched in unique immunologic signatures enriched in HIV-1-infected individuals and elite controllers. Parenthesis placed after the column name indicates the number of immune cell types counted in immunologic signatures in HIV-1-infected individuals and elite controllers. (E) Stacked bar chart representing the proportion of the different immune cell types in HIV-1-infected individuals before and after a short- and long period of ART. Statistic significance was calculated by Pearson’s chi-squared test by R with default options.

### Different combinations of immune cell signatures were observed alongside HIV-1 integrations associated with ART

In line with the finding that integration sites enrich specific immunologic signatures alongside HIV-1 infections associated with ART, I continuously investigated whether these enriched signatures can assign specific immune cells and proinflammatory soluble factors that states in the hypothesis of this work. I retrieved immunologic signatures uniquely enriched in pretreatment HIV-1-infected individuals (ut, 6 gene sets, 75%), HIV-1-infected individuals subjected to a short period of ART (st, 39 gene sets, 60%), HIV-1-infected individuals subjected to a long period of ART (lt, 128 gene sets, 84%), and elite controllers (15 gene sets, 68%). From these selected gene sets, I counted the appearance of names of immune cell types in the standard name of each enriched immunologic signature one by one (Figs 1D and 1E). I further classified all retrieved names of immune cells into 16 groups based on the Blood Cell Lineage chart published on the NIH website SEER Training Modules (Subtypes of cells in each cell category are described in Materials and Methods). It is important to note that ART interrupts HIV-1/AIDS progression; the composition of immune cell types observed here was in physiological HIV-1 infection conditions associated with ART. First, an increased variety of immune cell types in HIV-1-infected individuals subjected to a long period of ART (lt, found 132 terms of immune cells) was found (Fig 1D); among them, CD4 T cells (found 40 times, 30.3%) showed the highest frequency followed by myeloid cells (found 35 times, 26.5%). This composition of immune cell signature showed a difference between pretreatment HIV-1-infected individuals (ut, found 7 times) and patients subjected to a short period of ART (st, found 42 times) (Figs 1D and 1E, Pearson’s chi-squared test). The presence of the immune cell signatures in elite controllers (found 14 times) was more analogous to the latter cases, except for the group “B cells” (found 3 times, 21.4%) (Fig 1D). In contrast, the proportion of the group “B cells” showed a decrease from HIV-1-infected individuals subjected to a short-(found 7 times, 16.7%) to a long period of ART (found 9 times, 6.8%). The group “B cells” was not found in unique immunologic signatures enriched in pretreatment HIV-1-infected individuals (Fig 1E). Conversely, CD4 T cells, the main HIV-1 reservoir cell type, showed an increase of their proportion alongside HIV-1 infections associated with ART (ut, found 1 time, 14.3%; st, found 9 times, 21.4%; lt, found 40 times, 30.3%). A regression of the presence of CD8 T cells (ut, found 1 time, 14.3%; st, found 6 times, 14.3%; lt, found 11 times, 8.3%) was however observed after a long period of ART (Fig 1E). The proportion of myeloid cell signature remained relatively stable throughout infections (ut, found 2 times, 28.6%; st, found 15 times, 35.7%; lt, found 35 times, 26.5%) (Fig 1E). Intriguingly, the presence of NK cell signature was only observed in HIV-1-infected individuals subjected to a long period of ART (count 4 times, 3%) (Fig 1E). Collectively, these findings suggest that different combinations of immune cells with an increasing variety were present alongside HIV-1 infections with ART.

### Different combinations of proinflammatory soluble factor signatures were observed alongside HIV-1 integrations associated with ART

In the continuation of the previous logic, I also counted the appearance of the name of proinflammatory soluble factors, including cytokines and chemokines in each uniquely enriched immunologic signature (Fig 2A) and found that 1, 8, 33, and 6 unique immunologic signatures contained the name of proinflammatory soluble factors (Fig 2A). I further grouped these proinflammatory soluble factors based on their appearance alongside HIV-1 infections associated with ART: group I factors, including IL4 and IL12, were present in pretreatment HIV-1-infected individuals and throughout the whole infection process (Fig 2A); group II factors, including CXCR5, IFNG, IFNB, IL10, and TGFB were present in HIV-1-infected individuals subjected to ART irrespective of the period of ART (Fig 2A) and group III factors, including CXCL4, IFNA, IL1, IL2, IL6, IL7, IL15, IL18, TNF appeared only after a long period of ART (Fig 2A). To further investigate each group of proinflammatory soluble factors, I sought KEGG pathways [28-30] enriched by the genes retrieved from uniquely enriched immunologic signatures with corresponding proinflammatory soluble factors (Fig 2B). No clear separation of enriched KEGG pathways between HIV-1-infected individuals subjected to a short- and long period of ART was observed (Fig 2B), implying that the enrichment of these pathways was irrelevant to the period of the therapy. Nevertheless, enriched pathways in “cancer specific types” were often associated with IFNB (lt), CXCR5 (st), and IFNG (lt); enriched pathways in “signal transduction” and “immune system” were more related to IL10 (st) and IFNB (st); enriched pathways in “infectious disease viral” were more related to CXCR5 (st) (Fig 2B). As for group III factors, pathways in “signal transduction”, “immune system”, and “infectious disease viral” were prevalent for every proinflammatory soluble factor in this group (Fig 2C, the first column from the left). Pathways in “membrane transport” (hsaID uniq. IL12), “glycan biosynthesis” and “metabolism” (hsaID uniq. IL17), “carbohydrate metabolism” (hsaID uniq. CXCL4), “replication and repair” (hsaID uniq. CXCL4), and “endocrine and metabolic disease” (hsaID uniq. CXCL4) become specific to indicated proinflammatory soluble factors (Fig 2C). As I compared enriched KEGG pathways between group II and group III factors, more than half (61 pathways) of the pathways were in common (Fig 2D). Among them, “infectious disease viral”, “immune system” and “signal transduction” were the top three KEGG BRITE classifications containing enriched pathways in both groups of factors (Fig 2E). Enriched pathways in the classification “cancer specific types” were only found in group II factors (Fig 2E). Collectively, these findings suggest that different combinations of proinflammatory soluble factors participated in immunity alongside HIV-1 infections associated with ART.

**Fig 2.**
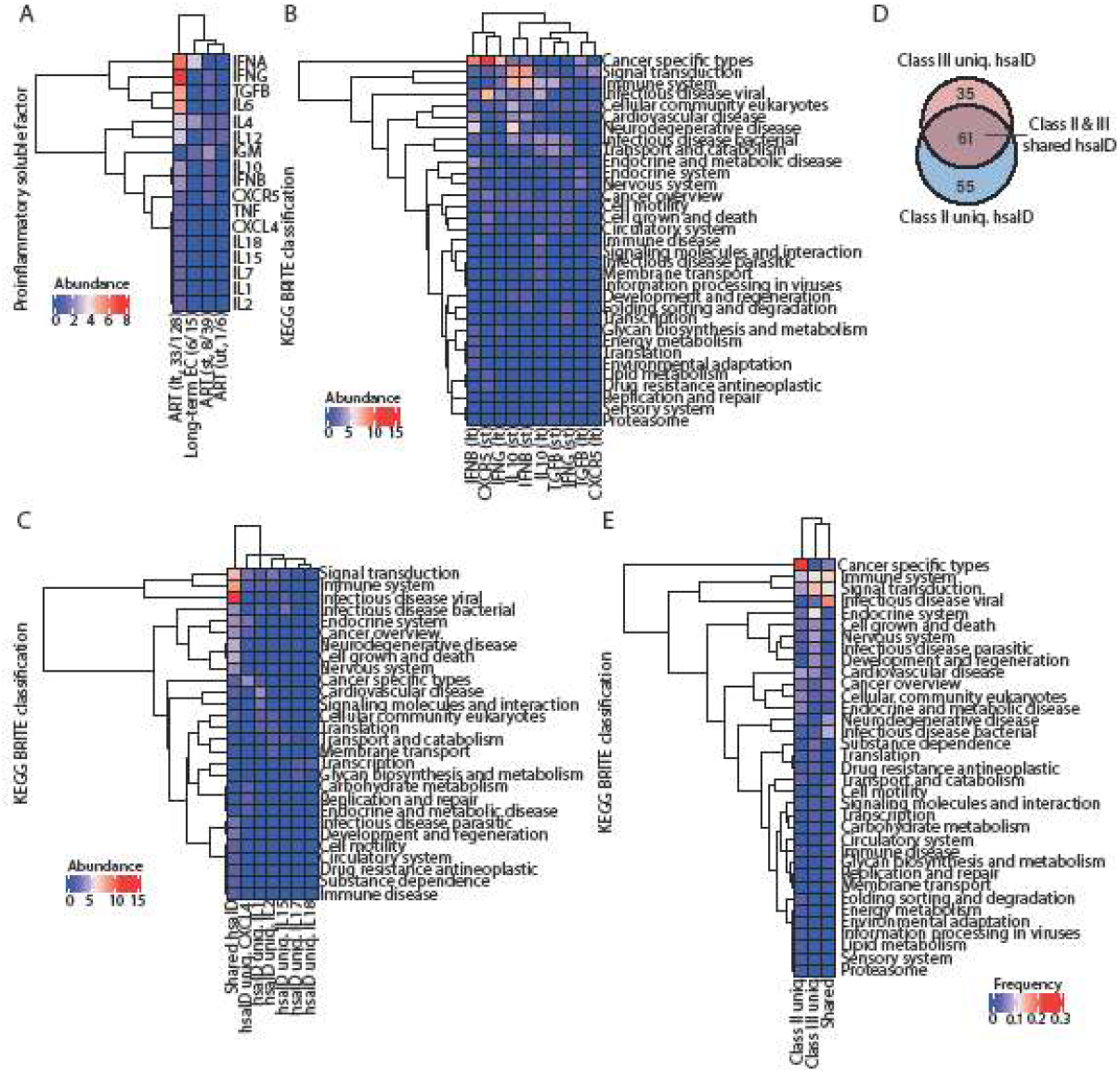
Different compositions of proinflammatory soluble factor signatures were present alongside HIV-1 infections associated with ART. (A) Cluster heatmap represents the abundance of proinflammatory soluble factors present in unique immunologic signatures enriched in HIV-1-infected individuals and elite controllers. Parenthesis placed after the column name indicates the quantity of immunologic signatures harboring proinflammatory soluble factors from total signatures in HIV-1-infected individuals and elite controllers. (B) Cluster heatmap representing the abundance of each KEGG BRITE classification harboring enriched pathways related to group II factors in HIV-1-infected individuals after a short-(st) and long (lt) period of ART. (C) Cluster heatmap representing the abundance of each KEGG BRITE classification harboring enriched pathways related to group III factors in HIV-1-infected individuals after a long period of ART. The abundance of KEGG BRITE classifications harboring enriched pathways present in all group III factors (so-called “Shared hsaID”) were compared to other enriched hsaID unique to every group III factor. (D) Venn diagram showing the overlap of enriched KEGG pathways represented by hsaID related to group II- and group III factors. (E) Cluster heatmap representing the frequency of shared- and unique hsaID enriched by gene sets related to group II- and group III factors. Each enriched KEGG pathway contains its own hsaID. Enriched pathways were classified in corresponding KEGG BRITE classifications.

To verify whether enriched pathways involved in “cancer specific type”, “immune system”, “signal transduction”, and “infectious disease viral” were truly required during HIV-1 infections, I withdrew all genes present in these four KEGG BRITE classifications from our input list and repeated the over-representation analysis performed in Figure 1A. First, I observed that the majority of genes present in “cancer specific types”, “immune system” and “infectious disease viral” are common, whereas the genes present in “signal transduction” appeared to be distinct from the other three classifications (Fig 3A). Intriguingly, genes present in these four KEGG BRITE classifications were restricted to HIV-1-infected individuals receiving ART (Figs 3B-3E), except the genes *rapgfe2* (Fig 3E) and *ppp3cc* (Figs 3C-3E), which were shown in pretreatment HIV-1-infected individuals as well. The gene *cyth1* was found in pretreatment HIV-1-infected individuals and patients subjected to a long period of ART; whether the absence of *cyth1* in HIV-1-infected individuals subjected to a short period of ART (Fig 3E) is due to technical reasons or it is a biological phenomenon will require further investigations. Furthermore, a decreased number of enriched immunologic signatures with minor significance (Fig 3G) was obtained as genes in these four KEGG BRITE classifications were withdrawn (Fig 3F) (S5-S12 Tables). And proinflammatory soluble factor signatures were infrequently present (Fig 3H), especially when the genes involved in “signal transduction” were taken away (Fig 3H, column 5). Consistently, a relatively weak correlation was present in the correlation plot as well (Fig 3J). However, signatures related to IL12 and IL18 were not affected (Fig 3H). The presence of the signatures relevant to IFNA and TGFB varied as genes in different KEGG BRITE classifications were withdrawn (Fig 3H, columns 1-5). The impact of the genes involved in these four KEGG BRITE classifications was minor in HIV-1-infected individuals subjected to a short period of ART (Fig 3H), thereby causing two clear clusters between a period of short- and long ART (Fig 3H). Nevertheless, a decrease in the correlation was still observed in HIV-1-infected individuals subjected to a short period of ART as genes in “immune system” were taken away (Fig 3I).

**Fig 3.**
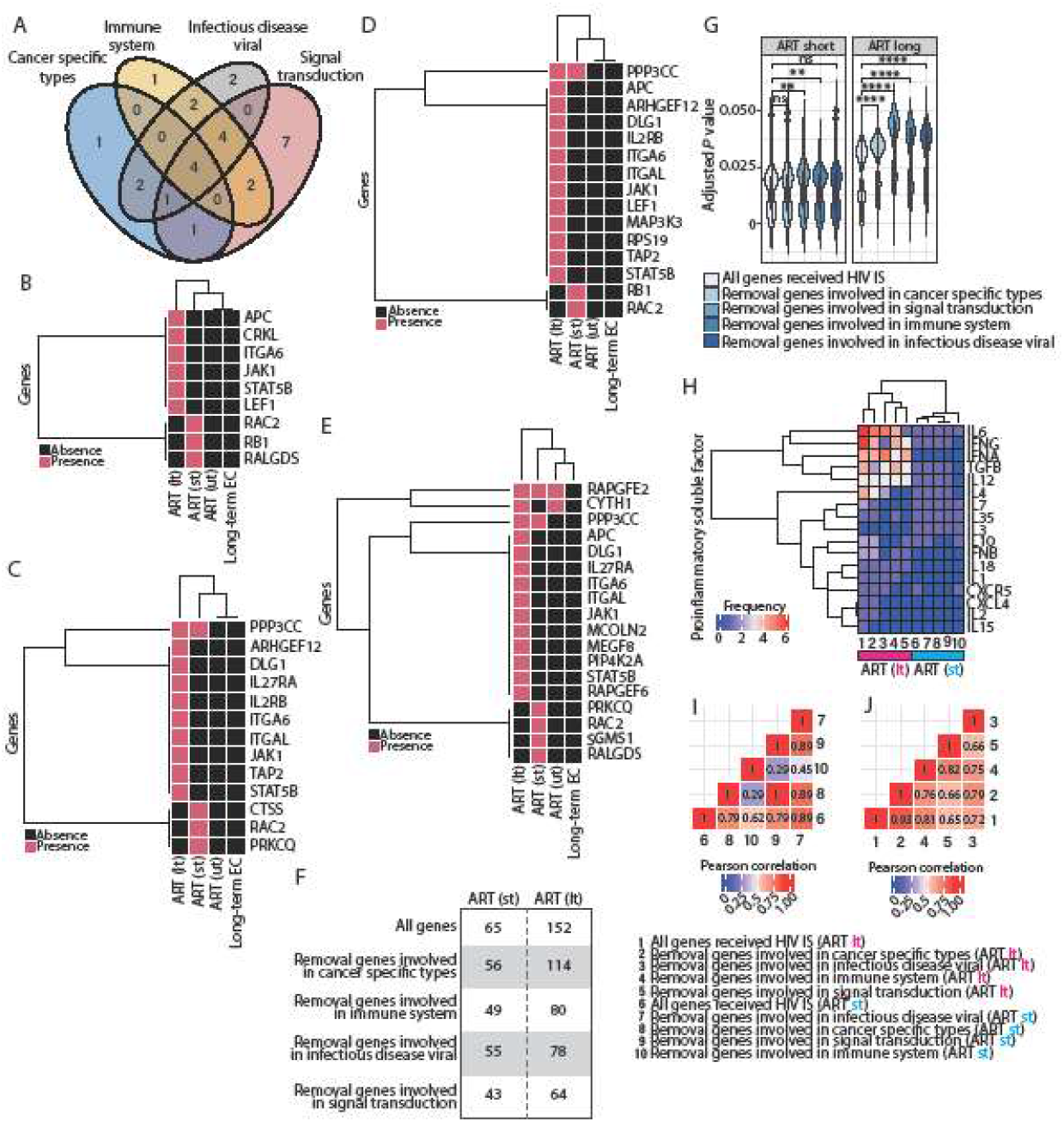
Top 4 KEGG BRITE classifications of the enriched pathways associated with enriched immunologic signatures in HIV-1-infected individuals subjected to ART. (A) Venn diagram representing the overlap of the genes present in enriched KEGG pathways involved in four KEGG BRITE classifications. (B - E) Cluster heatmap representing the presence of the genes present in enriched KEGG pathways involved in Cancer specific type (B), Immune system (C), Infectious disease viral (D), and Signal transduction (E) in HIV-1-infected individuals before ART (ut) and after a short- (st) or long period (lt) of ART and elite controllers. (F) Numbers of immunologic signatures enriched by either a complete list of the genes (All genes) or a gene list, from which the genes involved in these four KEGG BRITE classifications were withdrawn in HIV-1-infected individuals after a short- (st) or long period (lt) of ART. (G) Violin plots representing the distribution of adjusted *P* values corresponding to immunologic signatures enriched by either a complete list of the genes (All genes) or a gene list, from which the genes involved in these four KEGG BRITE classifications were withdrawn in HIV-1-infected individuals after a short- (st) or long period (lt) of ART. Statistic significance was calculated by the Wilcoxon test by R with default options. (H) Cluster heatmap representing the abundance of proinflammatory soluble factors present in each immunologic signature enriched by either a complete list of the (All genes, column 1 and column 6) or a gene list, from which the genes involved in selected four KEGG BRITE classifications were withdrawn in HIV-1-infected individuals after a short- (st, columns 2 - 5) or long period (lt, columns 7 - 10) of ART. (I - J) Correlation plots representing the Pearson correlation calculated based on the abundance of proinflammatory soluble factors in HIV-1-infected individuals after a short- (I) or long period (J) of ART.

### The majority of enriched immunologic signatures resulted from gene sets that interact with HIV-1

Numerous studies have highlighted the functions of several host genes, such as APOBEC (An et al., 2004), SAMHD1 (Laguette et al., 2011; Lahouassa et al., 2012), and TRIM5 (Javanbakht et al., 2006) are associated with HIV-1 pathogenesis and disease progression. I thus hypothesized that immunologic signatures and biological pathways enriched alongside HIV-1 infections may be conveyed by the host genes interacting with HIV-1. If this would be the case, I expect to observe that the enrichment is prone to be present in gene sets, that either show protein interactions with gene products of HIV-1 or affect HIV-1 replication and infectivity, whereas other gene sets that do not interact with HIV-1 are not responsible for the enrichment of any immunologic signature upon HIV-1 infections. To verify this hypothesis, I separated input genes reported to interact with HIV-1 from others that do not interact with HIV-1 (see Materials and Methods) and observed that 38.8% (47 genes out of 121 genes), 40.5% (60 genes out of 148 genes), 37.5% (66 genes out of 176 genes), and 29.8% (31 genes out of 104 genes) of the genes have been reported to interact with HIV-1. The majority of genes showed interactions with the HIV-1 *gag-pol* genes followed by the *tat* gene, the *env* gene, and the *nef* gene; few intact proviruses-targeted genes were reported to interact with the HIV *vpr, vif, rev*, and *vpu* genes (Fig 4A).

**Fig 4.**
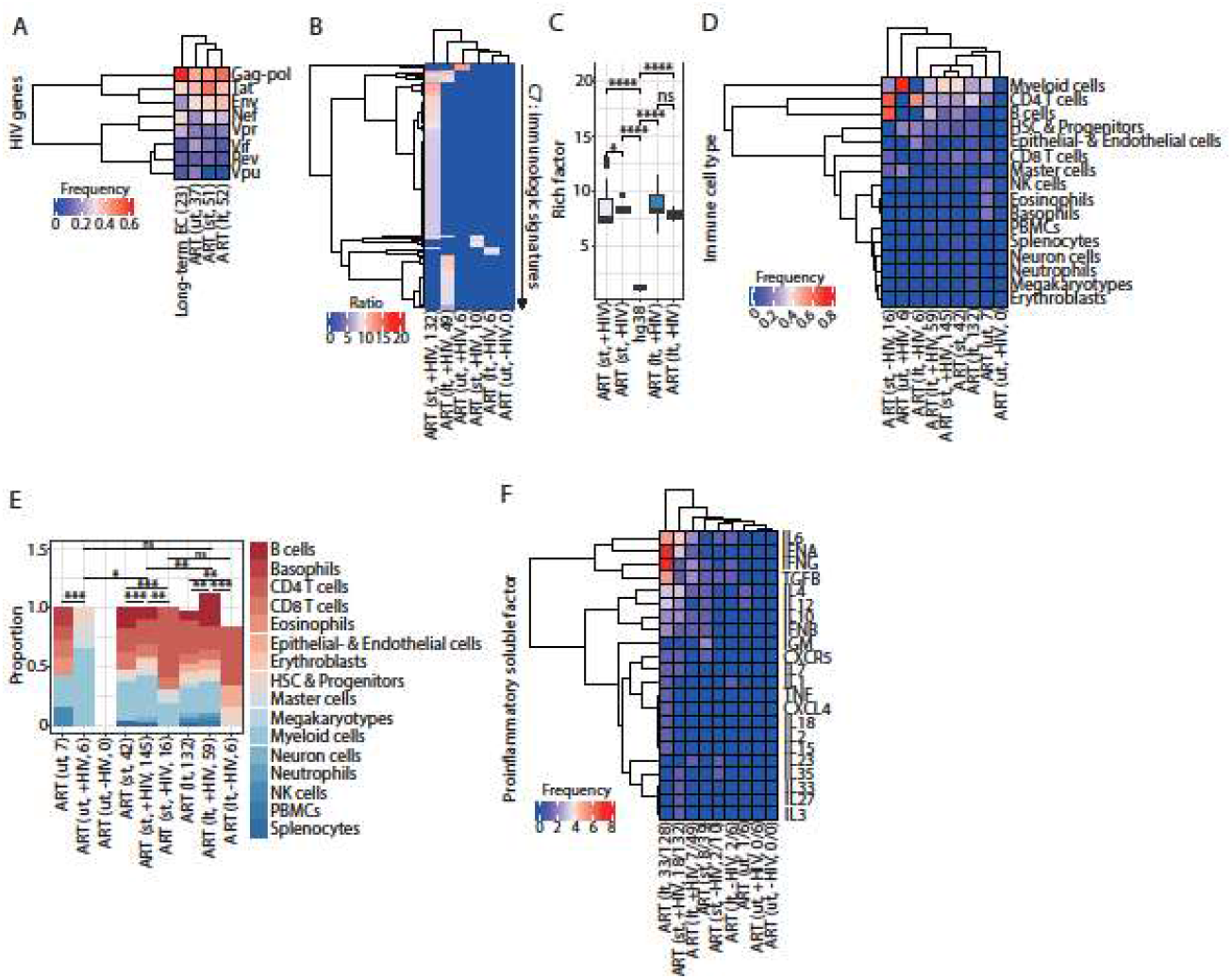
Intact provirus-targeted gene sets that interact with HIV-1 contributed to enrich immunologic signatures and pathways alongside HIV-1 infections associated with ART. (A) Cluster heatmap represents the frequency of intact proviruses-targeted genes interacting with HIV-1. Parenthesis placed after the column name indicates the total number of the intact proviruses-targeted genes reported to interact with HIV-1 based on the HIV-1 Human Interaction Database in HIV-1-infected individuals and elite controllers. (B) Cluster heatmap representing enriched immunologic signatures in HIV-1-infected individuals and elite controllers. Inside the parenthesis, +HIV and -HIV indicate intact proviruses-targeted genes were reported to interact or not, respectively; the number indicates immunologic signatures enriched in each scenario. The color scale represents the intensity of the enrichment represented by rich factors. (C) Box plots representing the magnitude of enrichment of immunological signatures. Commands executed to calculate rich factors were described in Materials and Methods. Statistic significance was calculated by the Wilcoxon test by R with default options. (D) Cluster heatmap representing the frequency of the immune cell types in immunologic signatures enriched in HIV-1-infected individuals. Inside the parenthesis, +HIV and -HIV indicate intact proviruses-targeted genes were reported to interact or not, respectively; the number indicates the quantity of immune cell types counted in each scenario. (E) Stacked bar chart representing the proportion of the different immune cell types present in enriched immunologic signatures in HIV-1-infected individuals before and after a short- and long period of ART. Statistic significance was calculated by Pearson’s chi-squared test by R with default options. Inside the parenthesis, +HIV and -HIV indicate intact proviruses-targeted genes were reported to interact or not, respectively; the number indicates the quantity of immune cell types counted in each scenario. (F) Cluster heatmap representing the abundance of proinflammatory soluble factors present in enriched immunologic signatures in HIV-1-infected individuals. Inside the parenthesis, +HIV and -HIV indicate intact proviruses-targeted genes were reported to interact or not, respectively; the number indicates the quantity of signatures harboring proinflammatory soluble factors out of the total number of enriched immunologic signatures.

I further performed the over-representation analysis on both subsets of input genes (interact with HIV-1 or not) and found that the enrichment of immunologic signatures was mainly contributed by the genes interacting with HIV-1 (Fig 4B) (S14 and S16 Tables). Only a tiny proportion of immunologic signatures was enriched by the genes, which were not reported to interact with HIV-1 (S15 and S17 Tables) (Fig 4B). Of note, genes interacting with HIV-1 in pretreatment HIV-1-infected individuals were only responsible for a small proportion of the enriched immunologic signatures; none of the immunologic signatures can be enriched using the genes, which were not reported to interact with HIV-1 in pretreatment HIV-1-infected individuals (Fig 4B) (S13 Table). Intriguingly, rich factors showed an increase in HIV-1-infected individuals subjected to ART no matter whether intact-provirus-targeted genes were reported to interact with HIV-1 or not (Fig 4C). Furthermore, immune cell types varied in enriched immunologic signatures using genes reported to interact with HIV-1 in patients receiving a short-(145 terms) and a long period (59 terms) of ART (Fig 4D-4E), resembling the similar circumstance using a complete list of input genes. In contrast, a poor combination of immune cell types was shown in enriched immunologic signatures using the genes, which were not reported to interact with HIV-1 (Fig 4D-4). Consistently, the majority of the proinflammatory soluble factors were present in enriched immunologic signatures using a complete list of input genes and the genes interacting with HIV-1 except in pretreatment HIV-1-infected individuals (Fig 4F). Either a tiny fraction or none of the proinflammatory soluble factor signatures were observed using the genes, which were not reported to interact with HIV-1 and using the genes retrieved in pretreatment HIV-1-infected individuals, respectively (Fig 4F). Altogether, these findings indicate that enriched immunologic signatures in HIV-1-infected individuals subjected to ART resulted from gene sets interacting with HIV-1.

## Discussion

In this work, I focused on integration sites of intact proviruses because they serve as a real latent reservoir and are responsible for a viral rebound (Ho et al., 2013). Based on the over-representation analysis of the genes targeted by intact proviruses, I first observed that different unique immunologic signatures were enriched alongside HIV-1 infections associated with ART (Fig 1A). Furthermore, an increase in variations of immune cell types was present in HIV-1-infected individuals subjected to a long period of ART (Fig 1D and 1E). Given that different kinds of innate immune-related cells are recruited to the site of HIV infection over time, unveiling the alterations of innate immune-related cells alongside HIV-1 infections can thus be essential to understand the evolution of HIV reservoirs. As shown in this work, various immune cell types were present in enriched immunologic signatures in HIV-1-infected individuals subjected to a long period of ART (Figs 1D and 1E), reflecting an environment with complicated immune responses. This observation of the variety of immune cell types can also link to my finding that showed abundant proinflammatory soluble factors in HIV-1-infected individuals receiving a long period of ART (Fig 2A). In addition to immune cells, it has also been known for a long time that infection with HIV-1 results in dysregulation of the cytokine profile (Clerici & Shearer, 1993), subsequently interfering with HIV-1 replication. This study also demonstrated that different compositions of proinflammatory soluble factor signatures were present at different stages of ART. I observed that CXCR5, IFNB, IFNG, IL10, and TGFB were found in enriched immunologic signatures in HIV-1-infected individuals subjected to a short- and long period of ART (Fig 2A); CXCL4, IFNA, IL1, IL2, IL6, IL7, IL15, IL18, and TNF were observed only in enriched immunologic signatures in HIV-1-infected individuals subjected to a long period of ART (Fig 2C). Notably, few proinflammatory soluble factors were observed in pretreatment HIV-1-infected individuals and elite controllers (Fig 2A). KEGG pathways enriched by gene sets related to group II- and III factors mainly fell into the KEGG BRITE classifications, “cancer-specific type”, “infectious disease viral”, “immune system” and “signal transduction” (Fig 2C). It is essential to note that gene sets from these four KEGG BRITE classifications were restricted to HIV-1-infected individuals subjected to a short- and a long period of ART, indicating that the selection of intact provirus integration can be disease progression-specific (Figs 3B-3E). This selection is irrelevant to elite control (Figs 3B-3E). Finally, I showed that enriched immunologic signatures were mainly contributed by genes interacting with HIV-1 (Fig 4B) even though the genes reported to interact with HIV-1 did not represent the majority of intact provirus-targeted genes in the input list. This observation may indicate that the affinity of the host genes towards gene products of HIV-1 can serve as a potential indicator to predict required immunologic signatures during HIV-1 infections associated with ART. At present, it is still not understood what makes the genes reported to interact with HIV-1 different from others; further investigations accompanied with more epigenetic features and a transcriptional status will be necessary.

Although it is presently workable to sequence the nearly full-length HIV-1 genome, the yield of retrieved sequences limits to the range of 50 to 100 sequences in cells purified from a patient (Hiener et al., 2017; Ho et al., 2013), thereby rendering accurate and quantitative investigation difficult. A better strategy to sequence the full-length HIV-1 genome of proviruses and elevate the yield of output sequences from clinical samples will be an absolute requisite shortly. On the other hand, it is thus fair to take into account a bias caused by a limited number of intact proviruses used in this work. Although enriched immunologic signature gene sets represented less than 10% of the input genes (S1-S4 Tables), rich factors showed significant enrichment (Fig 1B and Fig 4C), making results from the over-representation analysis trustable. A previous study also demonstrated that over-representation of pathways is not correlated to the number of genes in pathways (Chen et al., 2012). Several important questions will still need to be addressed: (1) whether host genes associated with innate immunity are more frequently to be targeted by intact proviruses. and (2) whether HIV-1 integration in genes relevant to innate immunity is due to transcriptionally active status, thereby being favorable for HIV-1 integration.

At present, transcriptional datasets that corresponded to input gene lists used in this study are not yet available. Therefore I employed another dataset, in which differentially expressed genes were computed from gene expression profiles from 30 HIV-1-positive- and 10 HIV-1-negative samples (Bai et al. 2022) and crossed this dataset with input gene lists. 3-(2.47% of input gene list from pretreatment HIV-1-infected individuals), 2- (1.34% of input gene list from HIV-1-infected individuals subjected to a short period of ART), 5- (2.84% of input gene list from HIV-1-infected individuals subjected to a long period of ART), and 6 genes (5.76% of input gene list from elite controllers) overlapped differentially expressed genes (S1 Fig), suggesting that the transcriptional status of a gene may also be involved in the interaction between HIV-1 integration sites and the host functional genome property. Further orthogonal approaches and additional transcriptomic datasets on corresponding patients’ cells will be requisites to clarify whether integration site frequency is dependent on levels of gene expression. Altogether, observations in this study allow opening a new avenue of research on HIV integration and the host functional genome property alongside HIV-1 infections associated with ART.

**S1 Fig.**
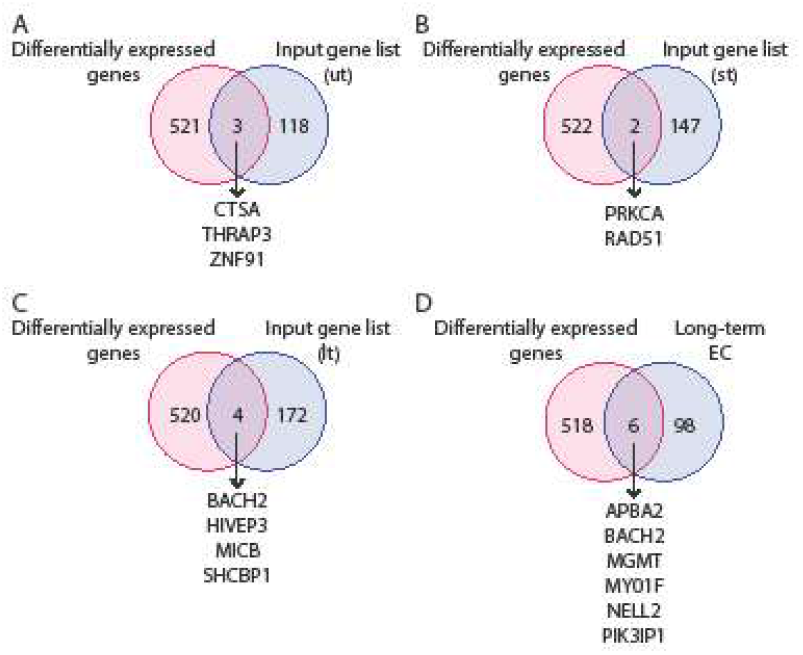
Overlaps between input gene lists and differentially expressed genes. Venn diagrams representing HIV-1-targeted genes that were observed in the differentially expressed gene list published by Bai et al. (2022) (Bai et al. 2022). 3-, 2-, 5-, and 6 genes showed an overlap with differentially expressed genes in pretreatment HIV-1-infected individuals (A), HIV-1-infected individuals subjected to a short period of ART (B), HIV-1-infected individuals subjected to a long period of ART (C) and elite controllers, respectively (D).

## Materials and Methods

### Acquisition and Procession of Public Datasets

Intact HIV-1 provirus integration sites were collected from Jiang et al. (2020) (Jiang et al., 2020) for elite controllers and Einkauf et al. (2022) (Einkauf et al., 2022) for pretreatment HIV-1-infected individuals and HIV-1-infected individuals subjected to a short- and long-period of ART. Integration sites retrieved from elite controllers were overlaid on Human_genome_GENCODE_v32.bed released from the GENCODE project (Frankish et al., 2019) to determine if the insertion of each integration site is intergenic or intragenic by exerting the command `intersect` with default options provided in bedtools (Quinlan & Hall, 2010). The list of 524 differentially expressed genes was retrieved from Bai et al. (2022) (Bai et al., 2022).

### Data and bioinformatic analyses MSigDb over-representation analysis

Enriched immunologic signatures were computed by employing the R package clusterProfiler (Version 4.4.1) (Wu et al., 2021; Yu et al., 2012) with the function enricher and default options. I performed the over-representation analysis (Boyle et al., 2004) using C7 immunologic signature gene sets curated in the Molecular Signatures Database (MSigDb) (Liberzon et al., 2011, 2015; Subramanian et al., 2005) as the background. Only the enriched immunological signatures with *p*-values (adjusted by the Benjamini-Hochberg method) smaller than 0.05 were selected. I further calculated a rich factor (Wu et al., 2021) to represent the enrichment score for every enriched immunologic signatures (Fig 1B) by dividing GeneRatio by BgRatio with command lines described below.

~~~
> MSigDb_output_file$GeneRatio <- **as.numeric**(**gsub**(“(\\d+)/(\\d+)”,
“\\1“, MSigDb_output_file$GeneRatio, perl=T)) /
**as.numeric**(**gsub**(“(\\d+)/(\\d+)”, “\\2“, MSigDb_output_file$GeneRatio,
perl=T))
*# Convert GeneRatio to numerical variables*.
> MSigDb_output_file$BgRatio <- **as.numeric**(**gsub**(“(\\d+)/(\\d+)”, “\\1“,
MSigDb_output_file$BgRatio, perl=T)) / **as.numeric**(**gsub**(“(\\d+)/(\\d+)”,
“\\2“, MSigDb_output_file$BgRatio, perl=T))
*# Convert BgRatio to numerical variables*.
> MSigDb_output_file <- MSigDb_output_file %>% **dplyr::mutate**(rich_factor
= GeneRatio/BgRatio)
*# Calculate rich factors*.
~~~

Of note, after performing MSigDB over-representation analysis, terms of immune cell types and proinflammatory soluble factors shown in the standard name of every enriched immunological signature gene set were retrieved for further statistical calculation. Based on the Blood Cell Lineage chart published on the NIH website SEER Training Modules (https://training.seer.cancer.gov/leukemia/anatomy/lineage.html), Immune cell types were classified into 16 groups, including (1) B cells, including plasma cells, (2) Basophils, (3) CD4 T cells, (4) CD8 T cells, (5) Eosinophils, (6) Epithelial- and Endothelial cells, (7) Erythroblasts, (8) Hematopoietic stem cells (HSC) & Progenitors, (9) Master cells, (10) Megakaryocytes, (11) Myeloid cells, (12) Neuron cells, including microglia, (13) Neutrophils, (14) NK cells, (15) Peripheral blood mononuclear cells (PBMC) and (16) Splenocytes in this study. A breakdown of the group (8) Hematopoietic stem cells (HSC) & Progenitors include stem cells and thymocytes; a breakdown of the group (11) Myeloid cells includes bone marrow-derived dendritic cells, bone-marrow-derived macrophage, dendritic cells, (monocyte-derived) macrophages, and macrophages. The frequency of the occurrence of immune cell types and proinflammatory soluble factors was represented as cluster heatmaps generated by employing the R package ComplexHeatmap (Gu et al., 2016) with default options. All plots present in this manuscript were generated in R with default options.

To obtain immunological signatures enriched by randomly selected genes and the whole genome, I retrieved protein-coding genes in humans from Human_genome_GENCODE_v32.bed released from the GENCODE project (Frankish et al., 2019) and either executed the R command `sample_n()` (with replacement) to randomly select 150, 250, 350, 450, 550, 650 and 1000 human genes or used all retrieved human genes to perform the MSigDb over-representation analysis described above.

### KEGG pathway over-representation analysis

Enriched KEGG pathways were retrieved by conducting KEGG pathways over-representation analysis built-in in the R package clusterProfiler (Yu et al., 2012) with the *p*-value cutoff equal to 0.5. Based on the hsaID, every enriched pathway was assigned to its corresponding KEGG BRITE classification to make the plots in Figure 2B-2D. Of note, to make Figure 2C, hsaIDs that were found in every group III factor were first separated from the hsaIDs that were unique to every single group III factor and then performed cluster heatmap on KEGG BRITE classifications between common and unique hsaIDs.

### Determination of intact-provirus-targeted genes interacting with HIV-1

In this study, I used the Human Interaction Database database to verify whether HIV-1-targeted genes interact with HIV-1 or not. This database documents all known interactions of HIV-1 and human genes that have been reported to affect HIV-1 replication and infectivity as well as interactions between HIV-1 gene products with host cell proteins, or proteins from disease organisms associated with HIV-1/AIDS (Fu et al., 2009; Pinney et al., 2009; Ptak et al., 2008).

### Statistics

All statistical tests were performed with R with default options. Details are provided where appropriate in the main text.

## Acknowledgments

I would like to thank Dr. Ariberto Fassati, Dr. Chien-Kuo Lee, and Dr. Zeger Debyser (alphabetic order) for their critical reading and constructive feedback on the manuscript.

