## Supplementary figures and images for "The dynamic linkage between intact provirus integration sites and the host functional genome property alongside HIV-1 infections associated with antiretroviral therapy"

### S1_Fig.pdf

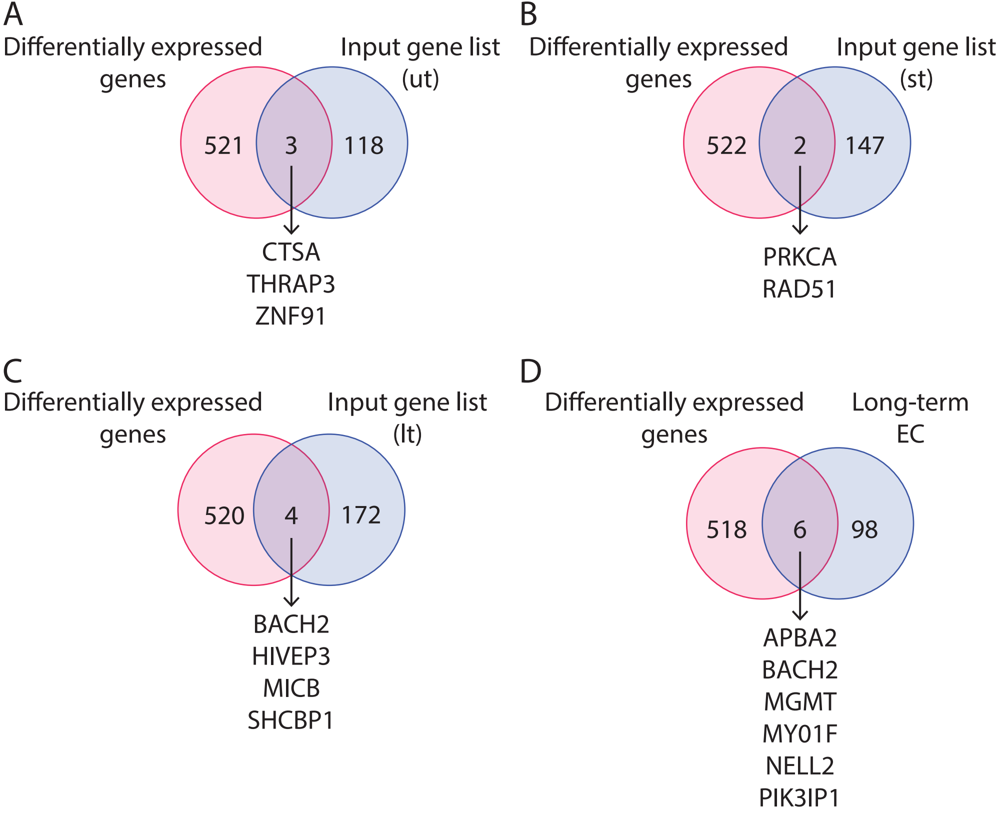
